# The N-end rule E3 ligase UBR2 activates Nlrp1b inflammasomes

**DOI:** 10.1101/429225

**Authors:** Hao Xu, Jianjin Shi, Zhenxiao Yang, Feng Shao, Na Dong

## Abstract

Innate immunity relies on the formation of different inflammasomes to initiate immune responses. The recognition of diverse infection and other danger signals by innate immune receptors trigger caspase-1 activation that induces pyroptosis. Anthrax lethal factor (LF) is a secreted bacterial protease that known to potently activate Nlrp1b inflammasomes in mouse macrophages, but the molecular mechanism underlying LF-induced Nlrp1b activation remains unknown. We here carried out both a mouse genome-wide siRNA screen and a CRISPR/Cas9 knockout screen seeking to identify genes that participate in Nlrp1b activation triggered by LF treatment. We found that the N-end rule pathway E3 ligase UBR2 is required for Nlrp1b activation and a ubiquitin conjugating E2 enzyme E2O is also involved in this process via its physically interaction with UBR2. We show that LF triggers activation of Nlrp1b by initiating the degradation of the N-terminal fragment of Nlrp1b itself that produced via an auto-cleavage process. This study deepens our understanding of innate immunity defense against bacterial infection by elucidating the functional role of UBR2-mediated N-end rule pathway in LF-induced Nlrp1b activation.

## Introduction

Vertebrates have evolved both innate and adaptive immune systems to fight infection, disease, and other unwanted biological invasions. The innate immune system is the first defense line; it recruits various classes of pattern-recognition receptors (PRRs) to coordinate rapid and versatile responses to these challenges^1^. PRRs, including Toll-like receptors (TLR), NOD-like receptors (NLR), and RIG-I-like receptors (RLR), sense pathogen-associated molecular patterns (PAMPs) or damage-associated molecular patterns (DAMPs), resulting in the release of pleiotropic effector molecules, reactive oxygen, nitrogen intermediates, costimulatory molecules, chemokines, and cytokines, etc. Various stimuli are recognized by specific PRRs, and signal transduction is subsequently propagated by the formation of a large protein platform known as inflammasomes, which activates caspase-1 protease. To date, there are five confirmed inflammasome components—distinguished based on their particular PRR protein—including NLRP1, NLRP3, NLRC4, AIM2, and pyrin inflammasomes^2–6^. The activation of caspase-1 leads to proteolytic processing of pro-IL-1β and pro-IL-18, which function as important ‘alarm’ cytokines to exaggerate immune responses to eliminate potential threats^7,8^. In addition, activated caspase-1 can cleave the pore-forming protein GSDMD, causing a massive release of cellular contents that is followed by a lytic cell death process known as pyroptosis^9–11^. Although great progress has been made in recent years towards understanding the complicated activation mechanism of inflammasomes, these processes remain unknown for several types of inflammasomes^5,12.^

NLRP1 was the first NLR reported to form an inflammasome complex^13^. Note that human have only one NLRP1 protein, but that the *Nlrp1* genes in mouse genome are highly polymorphic, encoding multiple paralogs of Nlrp1: Nlrp1a, 1b, and 1c. In addition to conserved domain including PYD, NACHT, and LRR, human NLRP1 has two additional domains at its C-terminus: a FIIND (function to find domain) and a CARD domain. Mouse Nlrp1 proteins lack the PYD^14^. The FIIND domain, which is structurally similar to ZU5-UPA autolytic proteolysis domain^15^, will undergo post-translational auto-cleavage, which is required for NLRP1 activity^15–17^. Following this proteolytic processing, the N-terminal and C-terminal cleavage products of NLRP1 remain associated in an auto-inhibited state, and C-terminal CARD of NLRP1 is able to recruit ASC and activate caspase-1 in over-expressed cells^16,17^. However, the activation mechanism of this unique from of NLR inflammation is still ambiguous.

It is well-established that LF can specifically activate Nlrp1b in mice and rats immune cells^18,19^; LF is an effector protein secreted by the virulent pathogen *Bacillus anthracis* during infection that functions biochemically as a metalloprotease enzyme which cleaves mitogen-activated protein kinase kinase (MEK) and thereby inhibits the MAPK signal transduction pathway^20,21^. However, MAPK inhibitors like U0126 and SB202190 did not affect Nlrp1b activation. Interestingly, only Nlrp1b variants from certain inbred mouse and rat strains are susceptible to LF-induced activation, for example the mouse BALB/c and 129/Sv strains are susceptible whereas the C57 BL/6J strain is not^19,22^. Studies have demonstrated that LF cleaves Nlrp1b at a position located after its N-terminal Lys44, and it is known that this cleavage is required for Nlrp1b activation^23,24,25^. But the precise mechanism underlying this is still poorly understood.

Nlrp1b activation can be specifically blocked by the 26S proteasome inhibitors MG132 and lactacystin^26,27^, while other inflammasomes are not affected. The N-end rule pathway —a protein-degradation system that exists in both eukaryotic and prokaryotic cells^28,29^—has been implicated in several studies to be involved in LF-triggered killing of macrophages. This protein-degradation system senses some specific N-terminal residues (N-degron) of peptides or proteins by UBR box-containing proteins (N-recognin) that function as E3 ligase to ubiquitinate N-degron for subsequent degradation. Further, preliminary data has indicated that amino acid derivatives can protect mouse macrophages from LF-triggered cell death^21,30^. However, the mechanisms through which Nlrp1b activation is linked to the N-end rule pathway and proteasomes is completely unknown.

To understand the mechanism of LF-induced Nlrp1b activation, we generated a RAW 264.7 mouse macrophage cell line stably expressing RFP-ASC and performed a mouse whole-genome siRNA screen using RFP-ASC aggregation as a phenotypic marker. We also carried out an unbiased genome-wide CRISPR-Cas9 genetic screen in immortalized BMDM cells to identify unknown components that function in LF-induced pyroptosis. Both screens identified the N-end rule E3 ligase UBR2 and further study confirmed that UBR2 is indeed required for LF-induced Nlrp1b activation. We also found that UBR2 function together with E2O in the N-end rule pathway-mediated Nlrp1b activation. Subsequent functional studies demonstrated that LF can trigger Nlrp1b activation through UBR2 mediated degradation of Nlrp1b N-terminal fragment produced by auto-cleavage. In total, our study shed new sights on the mechanism of LF-induced Nlrp1b activation and highlighted the important role of E3 ligase UBR2 in LF-induced N-end rule pathway-based degradation of Nlrp1b protein.

## Results

### The N-end rule pathway is involved in LF-induced Nlrp1b activation

To dissect the mechanism underlying Nlrp1b activation, we generated a mouse macrophage RAW 264.7 stable cell line expressing both RFP-ASC fusion protein and EGFP; this RAW^RA^ cell line is sensitive to anthrax LF. It is known that, during inflammasome activation triggered by bacterially expressed LF, the procaspase-1 p45 undergoes auto-cleavage to generate its active form p10, which subsequently lead to pyroptosis^22^. We found that in RAW^RA^ cell the auto-cleavage of procaspase-1 and cell death could be blocked by treatment with the 26S proteasome inhibitor MG132 and high concentration K^+^ ions (Figure 1A and S1A), findings consistent with previous reports^31,32^. Neither MG132 nor high concentration K^+^ disrupted the cleavage of MEK3 by LF, verifying intact protease activity of LF in treated cells. In RAW^RA^ cells incubated with LF, RFP-ASC aggregated within 1.5 h and the EGFP signal was completely absent by this point, owing to the release of intracellular contents during cell death. However, in the presence of MG132 or high concentration KCl, both RFP-ASC speck formation and cell death were blocked (Figure 1B). Live imaging revealed the dynamics of RAW^RA^ cells morphology changes as well as RFP-ASC speck formation induced by LF (Movie S1) and by the catalytically dead E687A mutant (Movie S2).

**Fig. 1.**
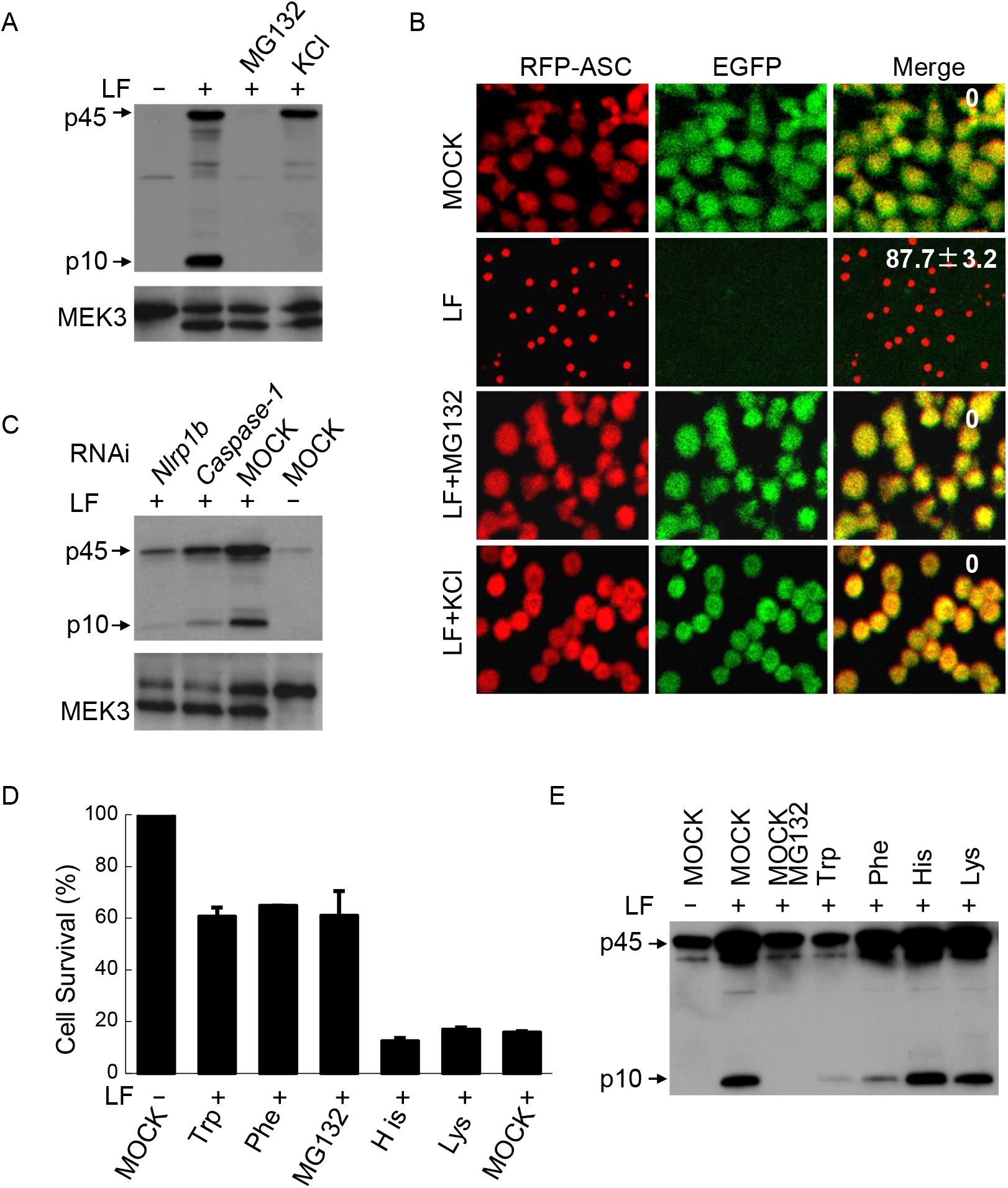
The N-end rule pathway is involved in LF induced Nlrp1b activation. (A) Effects of MG132 and KCl on caspase-1 activation in RAW^RA^ cells. RAW^RA^ cells were treated with LF WT (+) or E687C mutant (−) protein plus protective antigen (both 1 μg/ml) in the presence of 25 μM MG132 or 150 mM KCl as indicated. p10, mature caspase-1; p45, caspase-1 precursor. (B) RFP-ASC speck formation counting by confocal microscopy. RAW^RA^ cells were treated as in (A). The numbers in the merged panel are the mean value of the RAW^RA^ cells in which RFP-ASC aggregates formed. MOCK, addition of DMSO. (C) Effects of Nlrp1b and caspase-1 knockdown (4 mixed siRNAs) on inflammasome activation. Endogenous Nlrp1b or caspase-1 was silenced for 60 h before treatment with LF WT (+) or E687C mutant (−) protein plus protective antigen (both 1 μg/ml). MOCK, control scramble siRNA. (D) Inhibitory effects of basic and bulky hydrophobic amino acids on RAW^RA^ cells death triggered by LF. RAW^RA^ cells were pre-treated with 10 mM indicated amino acid before the addition of LF. Cell viability was measured using ATP cell viability assays. MOCK, DMDM medium. (E) Caspase-1 activation in the presence of different amino acid in *TLR4^-/-^* immortalized BMDM cells. RAW^RA^ cells were pre-treated with 10 mM indicated amino acid before addition of LF.

To determine whether the inflammasome activation and cell death that we observed in RAW^RA^ cells was mediated by functional Nlrp1b, the endogenous Nlrp1b was knocked down using siRNA, and these knockdown cells were treated with LF. As expected, knockdown of Nlrp1b significantly attenuated caspase-1 activation (Figure1C), formation of RFP-ASC speck, and pyroptosis in RAW^RA^ cells induced by LF (Figure S1B–C); there was no difference in the extent of MEK3 cleavage between LF-treated and control cells. We also found that caspase-1 knockdown impaired caspase-1 activation and cell death, although the formation of RFP-ASC speck was comparable to control cells (Figure 1C and S1B–C). Given that treatment of our newly generated RAW^RA^ stable cell line with MG132 or KCl as well as knockdown of Nlrp1b strongly inhibited the formation of RFP-ASC speck, the maturation of caspase-1, and the subsequent pyroptotic cell death, it is clear that this cell line is a suitable tool for use in monitoring LF-induced Nlrp1b activation.

Previous reports have implicated the N-end rule pathway in LF-induced Nlrp1b activation^30,33^. In this pathway, two specific types of N-terminal amino acids, referred to as N-degrons, are specifically recognized by E3 ubiquitin ligases that then catalyze ubiquitin chains on N-degron bearing target proteins, leading to their eventual degradation in the proteasome. Type 1 N-degrons bear N-terminal basic residues, (*e.g*., His, Lys, and Arg), while Type 2 N-degrons bear N-terminal bulky hydrophobic amino acids (*e.g*., Leu, Ile, Phe, Trp, and Tyr, etc). Guided by this knowledge, basic or bulky hydrophobic amino acids are often used experimentally as E3 ligase inhibitors for studying the N-end rule pathway^30^, and we thusly pretreated RAW^RA^ cells with different N-degrons for 30 minutes prior to LF delivery to determine whether the N-end rule pathway function in Nlrp1b activation. More than 60% of the RAW^RA^ cells that had been treated with bulky hydrophobic residues, Trp and Phe, survived the LF treatment, whereas, for both untreated and pre-treated with basic residues, fewer than 20% survived (Figure 1D). Additionally, immunoblotting revealed that Trp and Phe, but not His or Lys, strongly blocked the activation of caspase-1 in RAW^RA^ cells (Figure S2A). This data reveals inhibition of N-end rule pathway attenuates the activation of caspase-1 and subsequent pyroptosis.

To further confirm the role of the N-end rule pathway in Nlrp1b activation, we performed the same assay but used *Tlr4^-/-^* immortalized BMDM cells; these cells respond normally to LF treatment, but because they lack Tlr4, we were able to exclude the possibility of inflammasome activation by contaminated LPS present in recombinant LF and protective antigen proteins. Western blotting analysis showed that Trp and Phe, but not His or Lys, impeded the activation of caspase-1 in these cells (Figure 1E). Consistently the aforementioned data from RAW^RA^ cells, the formation of RFP-ASC speck was dramatically inhibited by Trp and Phe, but not His or Lys (Figure S2B and S2C). These data together confirm that the N-end rule signaling pathway is required for LF-induced Nlrp1b activation.

### The N-end rule E3 ligase UBR2 is required for Nlrp1b activation

To explore the underlying mechanism of LF-induced Nlrp1b activation, RAW^RA^ cells were used for a genetic screen based on a QIAGEN whole genome siRNA library. Considering that to date no unequivocal upstream signaling component that couples to Nlrp1b inflammasomes has been confirmed, we employed a “double check” method for the screen: this included both an RFP-ASC speck-forming assay and an ATP cell viability assay. We found that silencing of *UBR2*—encoding an E3 ligase involved in N-end rule signaling—lead to deficient Nlrp1b activation. The siRNA mixture of *UBR2* in QIAGEN library significant inhibited both RFP-ASC speck formation (Figure 2A) and cell death (data not shown). qPCR verified *UBR2* silencing (Figure S3A), and follow up experiments showed that two of the four siRNA targeting *UBR2* in the QIAGEN library showed apparent inhibitory effects on cell death (Figure S3B). We also carried out an unbiased genome-wide genetic screen using CRISPR-Cas9 technology to identify new components in LF-induced pyroptosis. The screen was carried out in immortalized *TLR4^-/-^* BMDM cells. As expected, *Antxr2, casp1, Nlrp1b* and *ASC* were among the most enriched genes. Moreover, *UBR2* was also strongly enriched. As the only E3 identified, two out of five *UBR2*-targeting gRNAs appeared in the top 30 hits (Fig. S3C). These results strongly indicated the direct role of the N-end rule pathway E3 UBR2 in LF-induced Nlrp1b activation.

To further validate the function of UBR2, two additional *UBR2* siRNAs and the *UBR2* mixture siRNA from the QIAGEN library were used to test caspase-1 activation in native 129/Sv mouse BMDMs. Compared with control scramble siRNA, the *UBR2* siRNAs and *Nlrp1b* siRNA caused remarkable decreases in the amount of capase-1 subunit p10 (Figure S4A), and qPCR revealed an approximate 30%-50% reduction of *UBR2* mRNA levels in the knockdown BMDMs (Figure S4B). Separate experiments in RAW^RA^ cells stably expressing *UBR2* shRNA (Figure S4C) showed that the both capase-1 and RFP-ASC speck-forming were almost completely blocked (Figure 2B and S4D), further verifying that UBR2 functions upstream of the Nlrp1b inflammasomes complex formation. We also successfully reconstituted Nlrp1b inflammasomes in human none-immune cell line 293T by expressing 129/Sv mouse derived Nlrp1b, human pro-caspase-1, and human pro-IL-1β (Figure S4E), and observed that the knockdown of endogenous *UBR2* in 293T cells also impaired LF-induced activation of the reconstituted inflammasomes (Figure 2C).

**Fig. 2.**
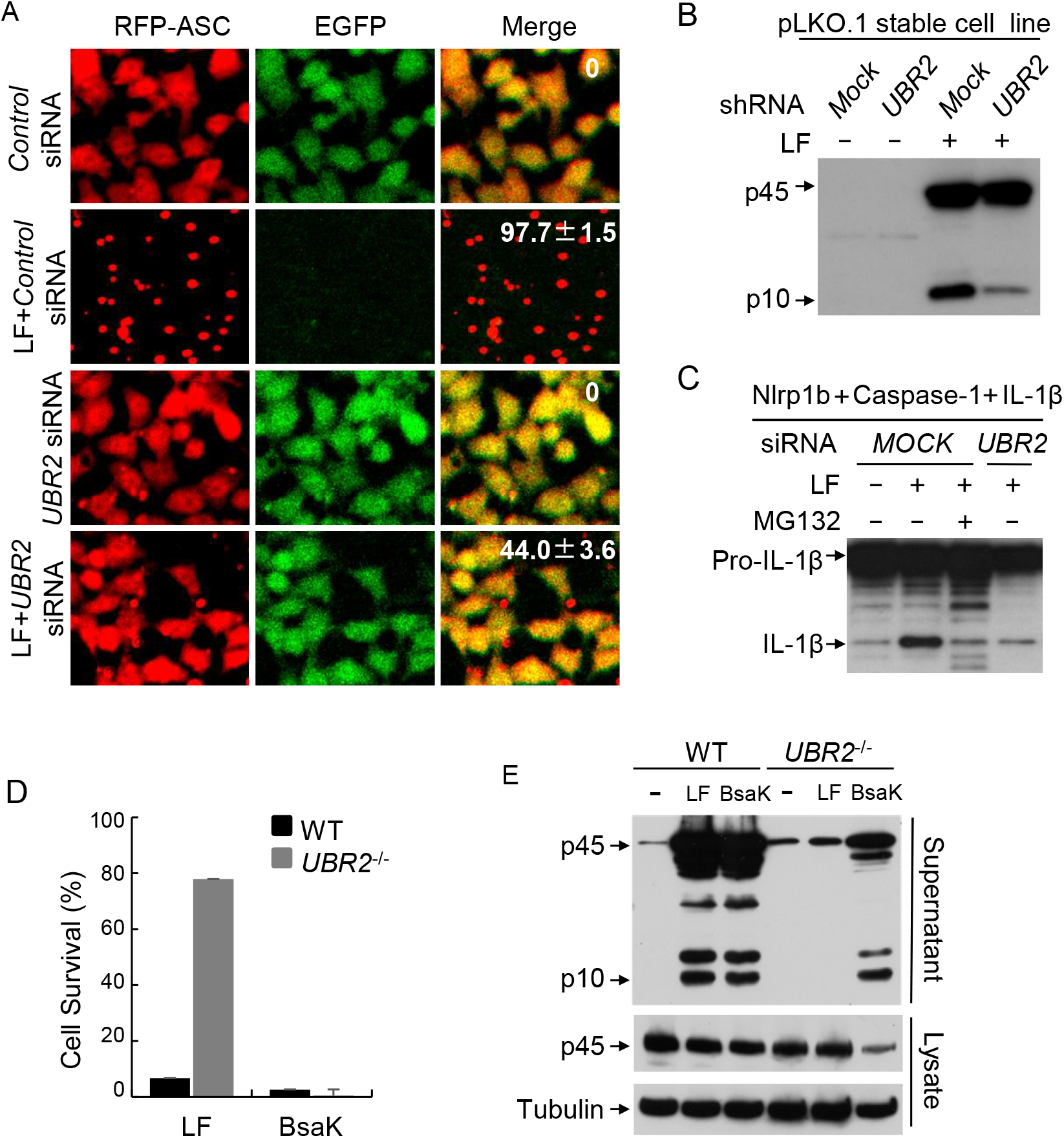
The E3 ligase UBR2 is required for LF-induced Nlrp1b activation. (A) RFP-ASC speck formation counting by confocal microscopy. RAW^RA^ cells were transfected with control siRNA or an *UBR2* siRNA mixture (QIAGEN library), for 60h followed by LF treatment. The numbers in the merged panel are the mean value of the RAW^RA^ cells in which RFP-ASC aggregates formed. (B) Effect of *UBR2* stable knockdown on capase-1 activation in RAW^RA^ cells. RAW^RA^ cells stably expressing *UBR2* or control scramble siRNA were incubated with LF WT (+) or E687C mutant (−) protein plus protective antigen (both 1 μg/ml) for 3h. MOCK, control scramble siRNA. (C) Knockdown of human *UBR2*(siRNA *UBR2*-05) inhibited the activation of reconstituted Nlrp1b inflammasomes in 293T cells. MOCK, control scramble siRNA; +, WT lethal toxin protein; –, lethal toxin E687C mutant protein. (D) Cell viability (percentage) in *UBR2*-knockout iBMDM cells. WT and UBR2^-/-^ cells were treatment with LF or BsaK plus protective antigen (both 1 μg/ml) for 3h. cell viability was measured using ATP cell viability assay. (E) Capase-1 activation in WT and *UBR2*-knockout iBMDM cells. WT and UBR2^-/-^ cells were incubated with LF or BsaK plus protective antigen (both 1 μg/ml) for 3h.

To further confirm the role of UBR2, immortalized BMDM *UBR2^-/-^* cells were generated using CRISPR-Cas9 mediated targeting (in the *Tlr4^-/-^* background). Knockout of *UBR2* in iBMDM cells dramatically blocked LF-triggered pyroptosis, while BsaK-induced cell death was not affected (Fig 2D). The absence of UBR2 also blocked caspase-1 processing triggered by LF treatment but not by BsaK treatment (Fig 2E). These results demonstrate that the N-end rule signaling pathway E3 ligase UBR2 is specifically required for LF-induced Nlrp1b activation.

### UBR2 functions together with E2O in the N-end rule pathway mediated Nlrp1b activation

UBR2 (ubiquitin protein ligase E3 component n-recognin 2) is a component of the N-end rule signaling pathway. Given that there is a total of 7 UBR box containing members (UBR1 to UBR7) encoded by mammalian genome, we were interested in testing whether other UBR E3 ligase members may function in Nlrp1b activation. RNAi-based experiments indicated that among the 7 *UBR*s, only *UBR2* (siRNA *UBR2*-05), was essential for Nlrp1b activation (Figure 3A). Note that the western blotting against MEK3 showed knockdown of *UBR*s did not affect the function of LF in cytosol.

**Fig. 3.**
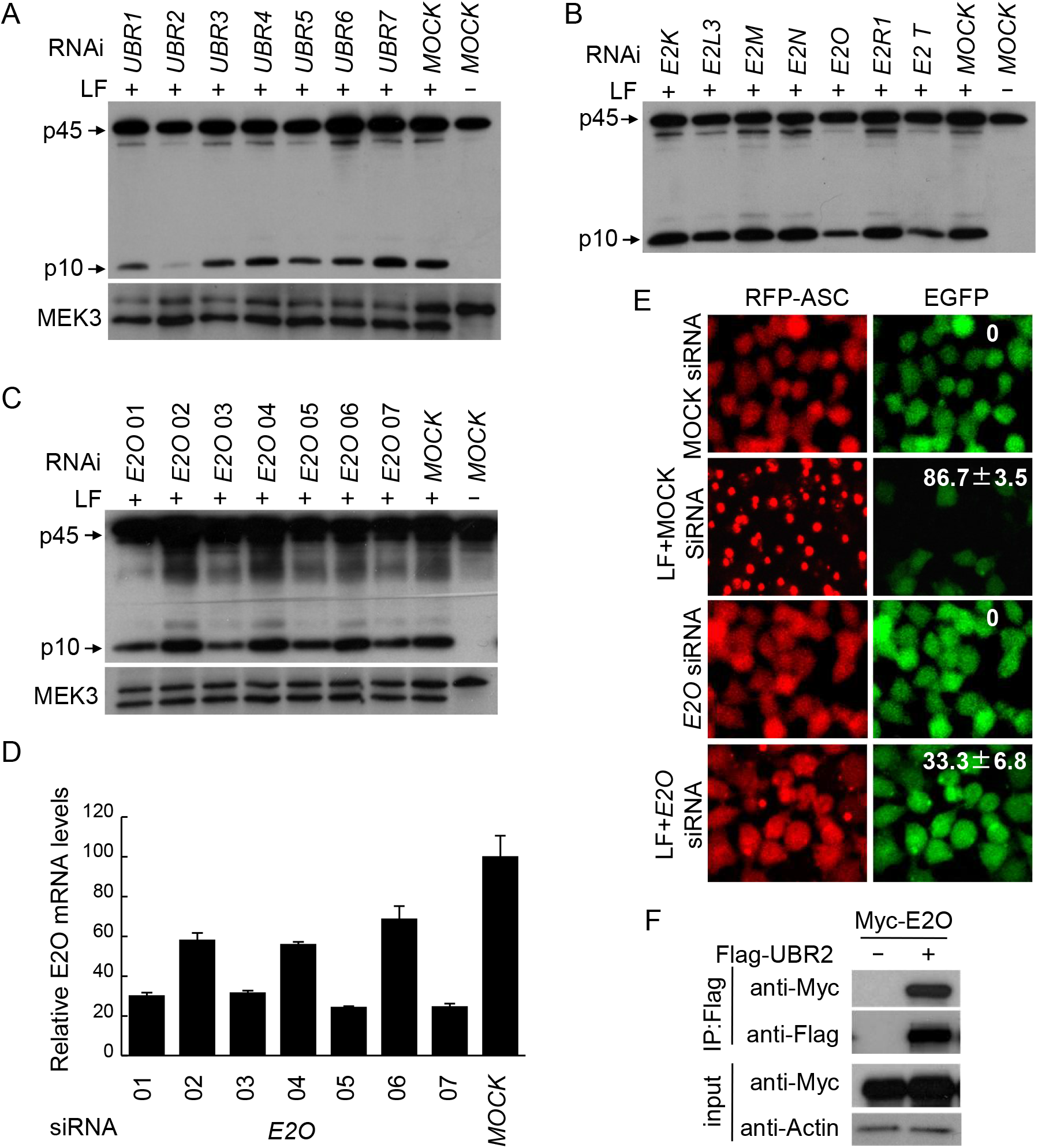
E2O functions together with UBR2 to regulates Nlrp1b activation. (A) Caspase-1 activation in *UBRs*-knockdown 129/Sv mouse BMDM cells. BMDM cells were transfected with the indicated siRNA for 60 h, then treated with WT LF (+) or E687C mutant (−) protein plus protective antigen (both 1 μg/ml) for 3h. Caspase-1 activation was validated by p10 immunoblotting. MOCK, control scramble siRNA. (B) Caspase-1 activation in E2s-knockdown 129/Sv mouse BMDM cells. BMDM cells were transfected with indicated siRNA for 60 h, then treated with lethal toxin (+) or E687C mutant (−) protein plus protective antigen (both 1 μg/ml) for 3h. Caspase-1 activation was validated by p10 immunoblotting. MOCK, control scramble siRNA. (C) Additional *E2O* siRNAs were used to confirm the inhibitory effects of *E2O* knockdown on caspase-1 activation. BMDM cells were transfected with the indicated siRNA respectively and then induced by LF plus protective antigen (both 1 μg/ml) for 3h. MOCK, control scramble siRNA. (D) *E2O* knockdown efficiency of samples in (C) measured by qPCR. (E) RFP-ASC speck formation counting in RAW^RA^ cells after knockdown of endogenous *E2O* by siRNA *E2O*-01. Statistics means percentage of RAW^RA^ cells in which RFP-ASC aggregates. MOCK, control scramble siRNA. (F) Interaction of E2O and UBR2. Myc-E2O and Flag-UBR2 were co-expressed in 293T cells for Flag-tag immunoprecipitation.

Since UBR2 is an E3 ligase that is essential for Nlrp1b activation and our experimental confirmation that the proteasome inhibitor MG132 can potently inhibit this activation, we want to identify the ubiquitin conjugating enzyme (*i.e*., E2 enzyme) that must function as the connection between the inflammasome activation and proteasomes. To this end, we conducted a small siRNAs screen for the targeted knock-down of 28 generally accepted *E2*s in BMDMs derived from 129/Sv mice. Western blotting for the caspase-1 active subunit p10 showed that knockdown of two *E2*s—*E2O* and *E2T*—suppressed LF-triggered caspase-1activation (Figure 3B and Figure S5A); neither of these *E2*s knockdown affected protease activity of LF by checking with western blotting against MEK3 (Figure S5B).

Additional siRNAs were designed to further examine if E2O and E2T have functions related to LF-induced Nlrp1b activation. Assays using multiple siRNAs against each of these two E2 proteins showed that knockdown of *E2O* (Figure 3C and 3D), but not *E2T* (Figure S5C and S5D), resulted in a significant decrease in the extent of caspase-1 activation, implying that E2O but not E2T function as an E2 ubiquitin conjugating enzyme during LF-induced Nlrp1b activation. Consistently, RFP-ASC speck formation was blocked when endogenous *E2O* was silenced by *E2O* siRNA mixture (Figure 3E). Further investigations based on immunoprecipitation analysis of extracts from 293T cells co-transfected with E2O and UBR2 showed that UBR2 can pulled down ectopically expressed E2O (Figure 3F). Taken together, these results show that the ubiquitin conjugating enzyme E2O functions together with E3 ligase UBR2 to regulate Nlrp1b activation.

### LF activate Nlrp1b through inducing the degradation of its N-terminal fragment produced by auto-cleavage

The N-end rule proteasomal degradation pathway is supposed to degrade a protein with negative role in Nlrp1b activation through specific E3 ligase UBR2. In the process of this study, we invariably observed an obvious decrease in Nlrp1b protein level after LF treatment (Figure S6A-S6C). Further, consider that LF cleavage of Nlrp1b is known to produce a neo-N-terminus (starting with Leu) that is a classic type 2 N-degron. We were thus interested in the possibility that the specific degradation of Nlrp1b itself through the N-end rule pathway may cause Nlrp1b activation.

Different from other NLRs, the Nlrp1 protein contains both a C-terminal CARD domain and a unique domain called FIIND. This FIIND domain must undergo auto-proteolysis prior to Nlrp1 activation (Figure 4A and 4B). We generated a Nlrp1b^V988D^ mutation that abolished this auto-cleavage^17^, which indeed abolish Nlrp1b activation as monitored via RFP-ASC speck formation (Figure S6D). Whereas LF treatment did not cause any change in the level of cleaved C-terminal fragments present in cells, it did cause significant reduction in the levels of both full-length Nlrp1b and N-terminal auto-cleavage product of Nlrp1b (Figure 4B left and S6C). However, for the Nlrp1b^V988D^ variant, we did not detect any N-terminal products, regardless of the LF-treatment status; moreover, the levels of the full-length Nlrp1b^V988D^ variant protein was not affected by LF treatment (Figure 4B right). These findings raised the interesting possibility that FIIND domain-mediated auto-cleavage of Nlrp1b may be an enabling prerequisite step that is required for LF-induced Nlrp1b activation.

**Fig. 4.**
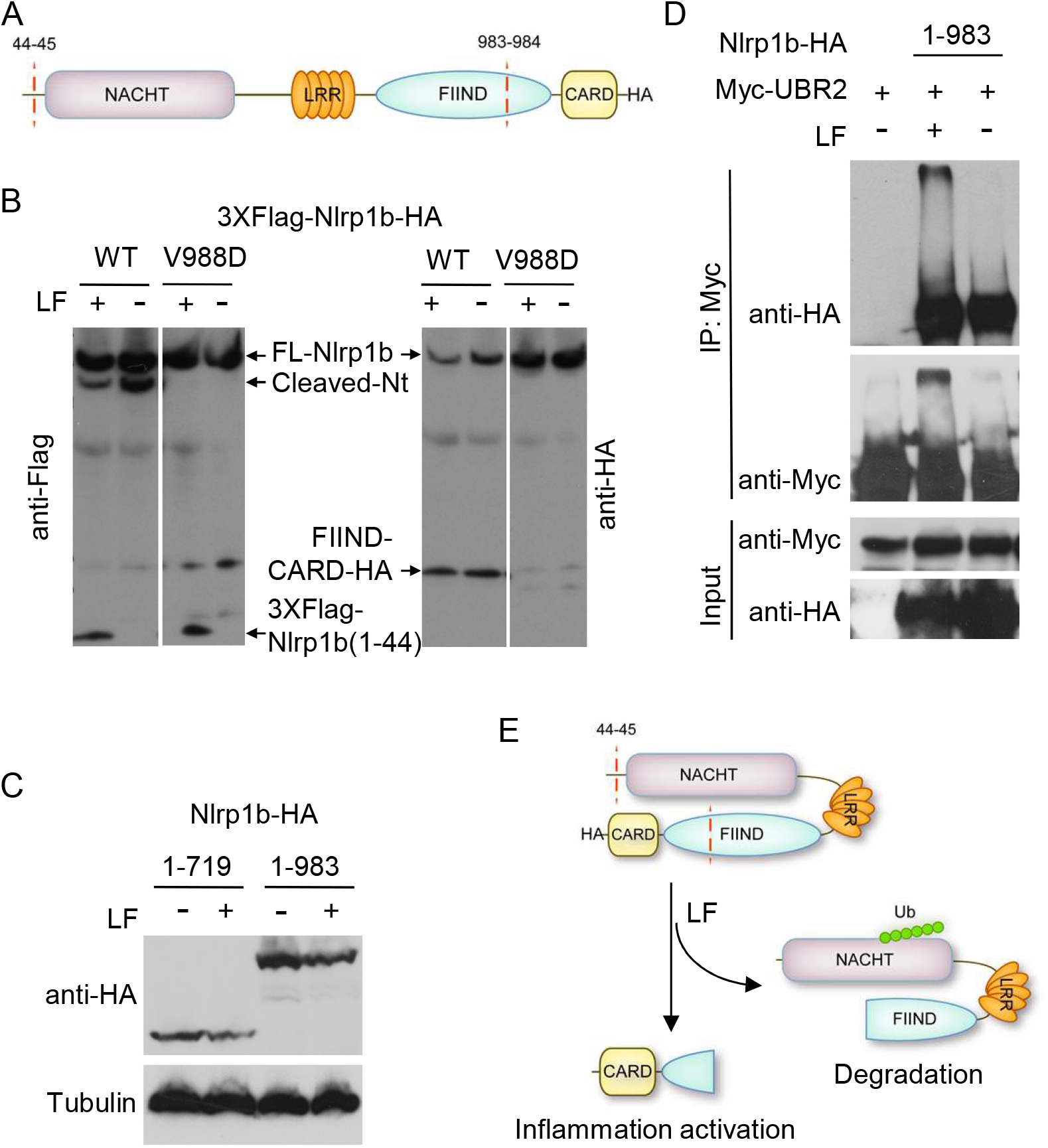
LF triggers UBR2 mediated Nlrp1b N-terminal degradation. (A) Domain structure of Nlrp1b. The LF cleavage site and FIIND auto-proteolysis site are labeled. (B) Expression of Nlrp1b and V988D mutant protein which lost the auto-cleavage ability. 293T cells were transfected with tagged Nlrp1b plasmids for 24 h, followed by induction with LF (+) or E687C mutant (−) protein plus protective antigen (both 1 μg/ml) for 3h. Nlrp1b and cleavage products after LF treatment were detected by Flag antibody. The C-terminal fragment after proteolytic cleavage within the FIIND domain was detected by HA antibody. (C) Effect of LF on protein expression of Nlrp1b1–719-HA or Nlrp1b1–983-HA. Cells were treated with or without LF plus protective antigen (both 1 μg/ml) for 4h after 24h of transfection. Protein expression were detected by western blotting with anti-HA and anti-Tubulin antibodies (representative of total protein expression). (D) Myc-UBR2 was were co-transfected with Nlrp1b1–719-HA or Nlrp1b1–983-HA into 293T cells for 24h. Then, LF plus protective antigen (both 1 μg/ml) were added for 4h before the cells were harvested for immunoprecipitation with anti-Myc magnetic beads. Protein expression was detected by western blotting with anti-Myc and anti-HA antibodies. (E) Proposed model of LF-induced Nlrp1b activation.

At this time, owing to a lack of a suitable antibody against the N-terminal FIIND domain auto-cleavage Nlrp1b product, we were not immediately able to conclusively determine whether the LF-treatment-induced decreases in the levels of both full-length and the N-terminal auto-cleavage product of Nlrp1b that we consistently observed are true reflections of these proteins in cells or are perhaps some artifact related to our Flag-antibody-based detection method. To discriminate between these two possibilities, we constructed several Nlrp1b that terminate before the FIIND cleavage site and that contain a C-terminal HA tag. Our observations that the levels of the Nlrp1b_1–719_-HA and Nlrp1b_1–983_-HA variant proteins were obviously reduced following LF treatment (Figure 4C) supported that the Nlrp1b N-terminal auto-cleavage produce may be a direct targeted of the N-end rule degradation pathway.

To explore this possibility, and recalling the requirement for UBR2 in LF-induced Nlrp1b activation, we co-transfected Nlrp1b_1–983_-HA with Myc-UBR2 and performed co-immunoprecipitation experiments with and without LF treatment. We found that UBR2 can interact with Nlrp1b_1–983_-HA truncation fragment, either with or without LF treatment, strongly suggesting that N-terminal of Nlrp1b can be targeted for ubiquitination by UBR2 to initiate the degradation process. Powerfully supporting this, we observed strong ubiquitination of Nlrp1b_1–983_-HA in cells treated with LF (Figure 4D). This result strongly demonstrates that LF can trigger the degradation of Nlrp1b itself through UBR2-mediated N-end rule pathway to activate Nlrp1b inflammasomes.

## Discussion

According to these data and previous studies^34^, we proposed a model of LF-induced Nlrp1b activation in which Nlrp1b’s autolytic cleavage C-terminal and N-terminal products that are present together interact in a manner that maintains an autoinhibition state. When LF is present, it triggers the cleavage and subsequent degradation of the N-terminal products—a process which the present study shows is mediated by the N-end rule pathway E3 ligase UBR2—thereby releasing the Nlrp1b C-terminal product from its inhibited state so that it can initiate the formation of the inflammasome. (Figure 4E).

The N-end rule pathway E3 ubiquitin ligase UBR2 recognizes N-degrons and LF is a metalloprotease that can produce type 2 N-degron, so our data reveal that the previously unknown link for LF-induced Nlrp1b activation and the function of N-end rule signaling pathway in this process is the proteasome degradation of the N-terminal product of Nlrp1b itself triggered by LF cleavage through UBR2 mediated N-end rule pathway. Genetic and biochemical data solved the puzzle existing for a long time and also clearly elucidates the importance role of UBR2 in Nlrp1b inflammation activation.

The ubiquitin conjugating enzyme E2O (also termed E2–230k), which was initially reported to function in rabbit reticulocytes^35–37^, is widely used for the determination of the N-end rule pathway due to its strong ubiquitin-dependent degradation activity. Here, our study show that E2O functions together with N-end rule pathway component UBR2 to regulate Nlrp1b activation. This finding lends further support to the conclusion that E2O does function directly in the N-end rule pathway. This is a significant finding, as to date the only confirmed E2 enzyme in this pathway is the ubiquitin conjugating enzyme HR6B (the mammalian homologue of yeast RAD6). Although we here present evidence that ectopically expressed E2O can interacts with UBR2, further experimental support will be required to confirm their physiological interaction and to precisely characterize the functional role of E2O in UBR2-mediated Nlrp1b activation.

Recent studies have established that DPP8/9 inhibitors can activate Nlrp1b inflammasomes and have shown proteasome inhibitors can block this activation^38,39^. There are five alleles of Nlrp1b. DPP8/9 inhibitors activate several subsets of Nlrp1b that is not the same as LF. Murine macrophages that express either allele 1 or 5 are susceptible to LF-mediated pyroptosis, but those that express allele 2, 3, or 4 are resistant, while macrophages containing Nlrp1b alleles 1, 2, and 3, but not alleles 4 and 5, were sensitive to DPP8/9 inhibitors. However, both LF- and DPP8/9 inhibitors-induced activations are known to require the FIIND domain auto-proteolysis. It will be interesting to experimentally test whether DPP8/9 inhibitors have the ability to activate human NLRP1 inflammasomes and to determine if DPP8/9 functions in NLRP1-dependent pyroptosis. To date, there are no confirmed activators of human NLRP1, a protein that is not susceptible to LF treatment because it lacks an LF cleavage site. The biggest difference in the NLRP1 domain architecture between human and non-primate mammalian orthologs is the existence of a PYD domain at the N-terminus of human NLRP1. Mutations of PYD are known to cause several skin disorders^40^, underscoring the importance of this domain in NLRP1 inflammasome activity. Further investigations will be required to unravel the complicated endogenous regulation of NLRP1 and characterize its detailed mechanism of activation in innate immunity.

## Acknowledgments

We thank Dr. Stephen Leppla for providing anthrax LF and protective antigen. We thank Liyan Hu for drawing the model. We also thank all the members in Shao lab for helpful discussions and kind assistance. This work was supported by grants from China NSFC (81788101), National Key Research and Development Program of China (2017YFA0505900 and 2016YFA0501500), the Chinese Academy of Sciences (XDB08020202) and the Austrian Science Fund (P-28915) to F.S, and China NSFC grants to N.D (81671987).

## Materials and Methods

### Antibodies and reagents

Antibodies for caspase-1 was obtained from Santa Cruz Biotechnology (SC-515) and AdipoGen (AG-20B-0044). Anti-Myc-tag mAb-Magnetic Beads (M047–11) was purchased from MBL. The other antibodies used in this study were HA (Covance), Flag M2, Myc and tubulin (Sigma-Aldrich), MEK3 (Cell Signaling), and UBR2 (Abcam). Homemade anti-Nlrp1b antibody was raised against the Nlrp1b N-terminal sequence (amino acids

4–20). Cell culture products were from Life technologies; all other chemicals were from Sigma-Aldrich unless noted otherwise.

### ATP cell viability assay and RFP-ASC speck forming assay

For assays without siRNA silencing, 3 × 10^5^ RAW^RA^ cells were plated into 24-well dishes 12 h before LF treatment. Cells were washed three times with serum free Dulbecco’s Modified Eagle Medium (DMEM, GIBCO), followed by addition of 0.5 ml serum free DMEM into each well with or without LF (1 μg/ml) plus protective antigen (1 μg/ml). If required, KCl (150 mM, Sigma) or MG132 (25 μM, Sigma) was added concomitantly with LF; Cells and culture medium were collected respectively 1.5 h later for ATP cell viability assay (Promega) and capase-1 p10 western blotting. RFP-ASC speck forming was monitored by confocal microscopy (Zeiss LSM 510 Meta Confocal). To assay the effects of the N-end rule pathway on Nlrp1b inflammasome activation, RAW^RA^ Cells were pretreated with four different amino acids (10 mM each, Sigma) for 30 min before addition of LF.

### Reconstitution of Nlrp1b Inflammasomes in 293T cells

To reconstitute Nlrp1b inflammasome, 129/Sv mouse derived Nlrp1b, human pro-caspase-1 and human pro-IL-1β expression plasmids were co-transfected in 293T cells plated into 6-well dishes. About 20 h after transfection, 293T cells were treated with LF WT or E687C mutant recombinant protein plus protective antigen (both 1 μg/ml) for 6 h in the presence or absence of 25 μM MG132. Cell lysates were made using lysis buffer containing 50 mM Tris, 150 mM NaCl, 1% Triton X-100, and protease inhibitor cocktail (Roche), pH 7.5. We monitored pro-IL-1β process via immunostaining of the cell lysates with the IL-1β antibody. If *UBR2* RNAi was required, 2 × 10^5^ 293T cells were plated into 12-well dishes and then transfected with 60 pmol siRNA using lipofectamine 2000 (Invitrogen). 24h after transfection, cells from each of the wells were resuspended and then divided equally and re-plated into 3 wells in 12-well dish; this was followed by another round of siRNA transfection about 12 h later. 24 h after the second round of siRNA transfection, cells were co-transfected with expression plasmids for Nlrp1b, pro-caspase-1, and pro-IL-1β. 20 h after transfection, cells were incubated with LF for 6 h for activation of pro-IL-1β cleavage. Maturation of pro-IL-1β was monitored as indicated above.

### RNAi screen in RAW^RA^ cells and BMDM cells

A mouse genome-wide RNAi screen (QIAGEN) using RAW^RA^ cells was carried out in 96-well plates. E2 screen was performed using BMDM cells with siRNA against 28 E2 family genes. Briefly, 2 × 10^5^ differentiated 129/Sv mouse BMDM cells were plated into 24-well dishes 24 h before RNA interference. 40 pmol siRNAs were used to knockdown endogenous E2 in BMDM cells with transfection kit INTERFERin (Polyplus). 60 h after siRNA transfection, Cells were washed three times with pre-warmed serum free DMEM and incubated with highly purified LF plus protective antigen (both 1 μg/ml, Dr. Stephen Leppla) for 3 h, followed by collection of medium for trichloroacetic acid protein precipitation and p10 western blotting.

### Gene knockdown by RNAi

For the knockdown of endogenous genes, 2 × 10^4^ RAW^RA^ cells were plated into 24-well dishes 12 h before siRNA transfection. siRNAs (20 pmol each, GenePharma or Dharmacon) were transfected using INTERFERin (Polyplus) following the product instructions. The ATP cell viability assays, RFP-ASC speck forming assays, and determination of caspase-1 activation were performed 60 h after siRNA transfection.

### Establishment of the stable shRNA knockdown cell line

A pLKO.1 plasmid was constructed via the insertion of a hairpin shRNA containing corresponding DNA of for the *UBR2*-05 siRNA listed in Figure S7. Scramble or stable knockdown of *UBR2* RAW^RA^ cell line was conducted following a previously developed protocol^41, 42^, and puromycin (2 μg/ml) was used for selection. The efficiency of *UBR2* knockdown was measured using qPCR, and the effects of *UBR2* knockdown on caspase-1 activation and RFP-ASC speck forming were assayed as indicated above.

### Genome-wide CRISPR-Cas9 screens

*Tlr4^-/-^* iBMDM cells stably expressing the Cas9 protein were seeded in the 15-cm dishes (2×10^6^ cells in 20 ml media per dish), and a total of 2×10^7^ cells were infected with a gRNA lentivirus library. 60 hours after infection, cells were re-seeded at a density of 1×10^5^ ml^-1^ in fresh media supplemented with 5 μg ml^-1^ puromycin to eliminate non-infected cells. After 6 to 8 days, 3×10^8^ cells from five culture dishes were treated with LF to trigger Nlrp1b-mediated pyroptosis, while another 3×10^8^ cells were left untreated as the control samples. Surviving cells were collected after reaching near 90% confluence and were lysed in the SNET buffer (20 mM Tris-HCl (pH 8.0), 5 mM EDTA, 400 mM NaCl, 400 μg ml^-1^ proteinase K, and 1% SDS). Genomic DNA were isolated using phenol-chloroform extraction and isopropanol precipitation method. The DNA was dissolved in water (4–5 μg μl^-1^), and was subsequently used as the templates for amplification of gRNA^11^.

### Generation of CRISPR-Cas9 mediated UBR2 knockout cell lines

Human codon-optimized Cas9 (hCas9) and GFP-targeting gRNA-expressing plasmids (gRNA_GFP-T1) were purchased from Addgene. The 19-bp GFP-targeting sequence in the gRNA vector was replaced with the sequence targeting the desired gene via QuickChange site-directed mutagenesis (stratagene). The sequence used to target mouse *UBR2* was GCGTCGGAGATGGAGCCCG. To construct the knockout cell lines, 1 μg of gRNA-expressing plasmid, 3 μg of hCas9 plasmid and 1 μg of pEGFP-C1 vector were co-transfected into iBMDM cells. Three days later, GFP-positive cells were sorted into single clones via flow cytometry using the BD Biosciences FACSAria II or the Beckman Coulter MoFlo XDP cell sorter and plated into 96-well dishes. Single clones were screened using T7 endonuclease I-cutting assay, and the candidate knockout clones were verified by sequencing of the PCR fragments and by western blotting^43^.

### Live Images

2 × 10^4^ RAW^RA^ Cells were plated into 96-well dishes 12 h before LF treatment. DIC, EGFP, and RFP images using a taken using PerkinElmer Ultra-View spinning disc confocal microscope.

---

**Figure S1.**
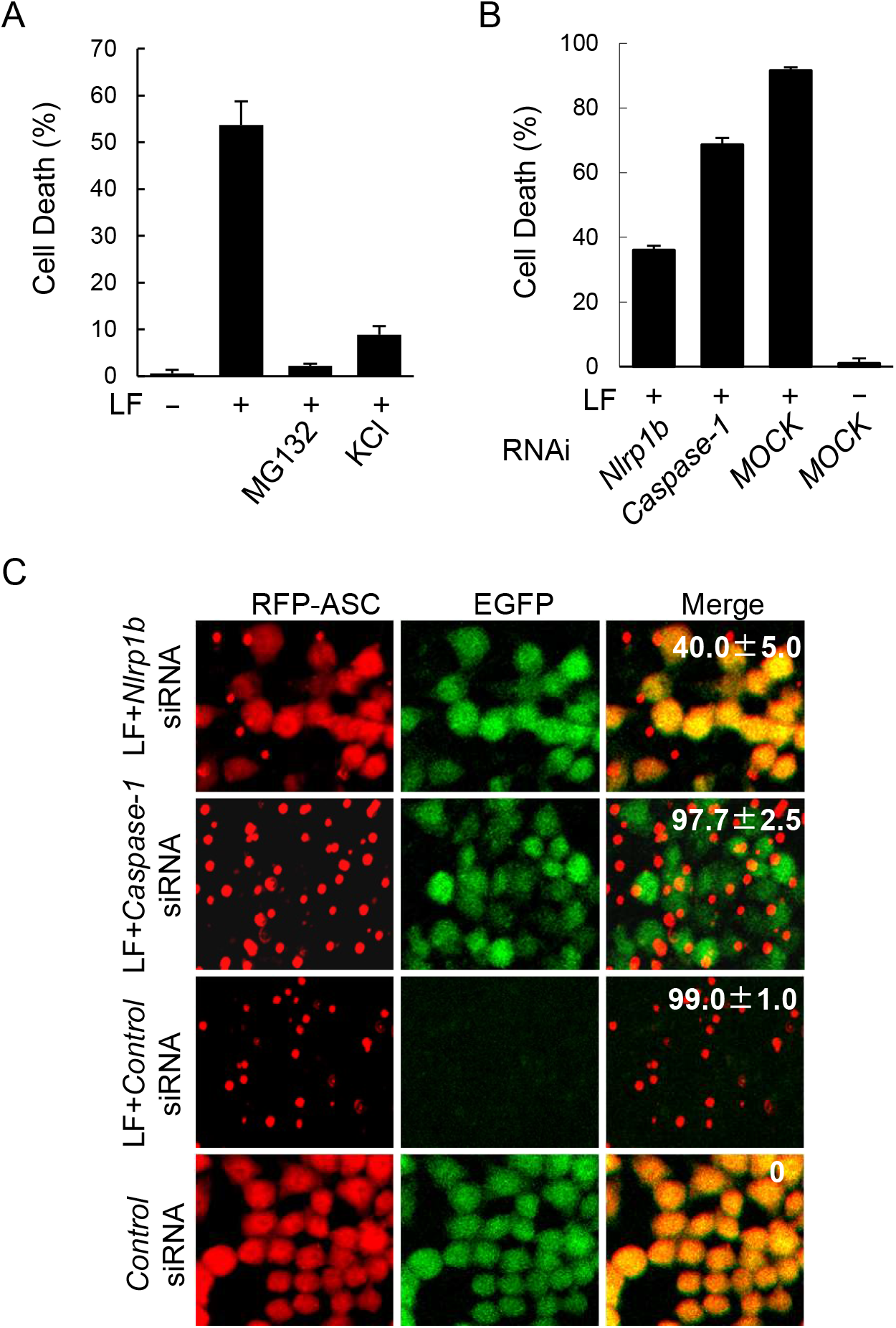
Establishment of RAW^RA^ stable cell line for the Nlrp1b inflammasome study. (A) Cell death measured by ATP cell viability assay. RAW^RA^ cells were treated with LF WT (+) or E687C mutant (−) protein plus protective antigen (both 1 μg/ml) in the presence of 25 μM MG132 or 150 mM KCl as indicated. Error bars mean standard deviations. (B) Effects of Nlrp1b and caspase-1 knockdown on cell death measured by ATP cell viability assay. Endogenous Nlrp1b or caspase-1 were silenced for 60 h before LF treatment. (C) RFP-ASC specks formation counting by confocal microscopy. RAW^RA^ cells were transfected with indicated siRNA for 60 h followed by LF treatment. Statistics means percentage of RAW^RA^ cells in which RFP-ASC aggregates.

**FigureS2.**
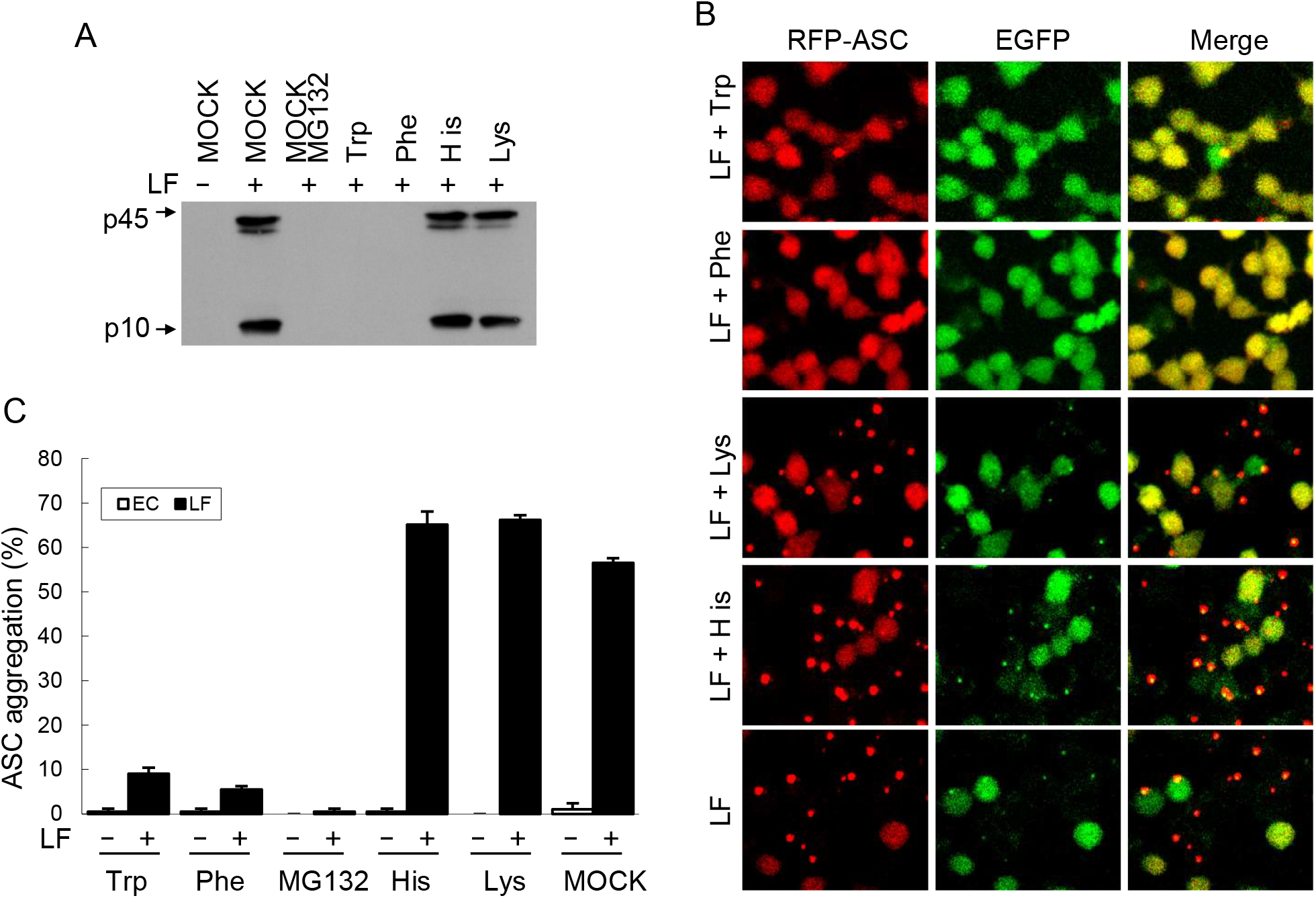
The N-end Rule pathway controls LF-induced Nlrp1b activation. (A) Inhibitory effects of basic and bulky hydrophobic amino acids on RAW^RA^ cells death triggered by LF. RAW^RA^ cells were pretreated with 10 mM indicated amino acid before addition of LF. (B) RFP-ASC specks formation in RAW^RA^ cells pretreated with indicated amino acids. MOCK, DMDM medium. (C) Statistics of RFP-ASC aggregation assayed in (B).

**Figure S3.**
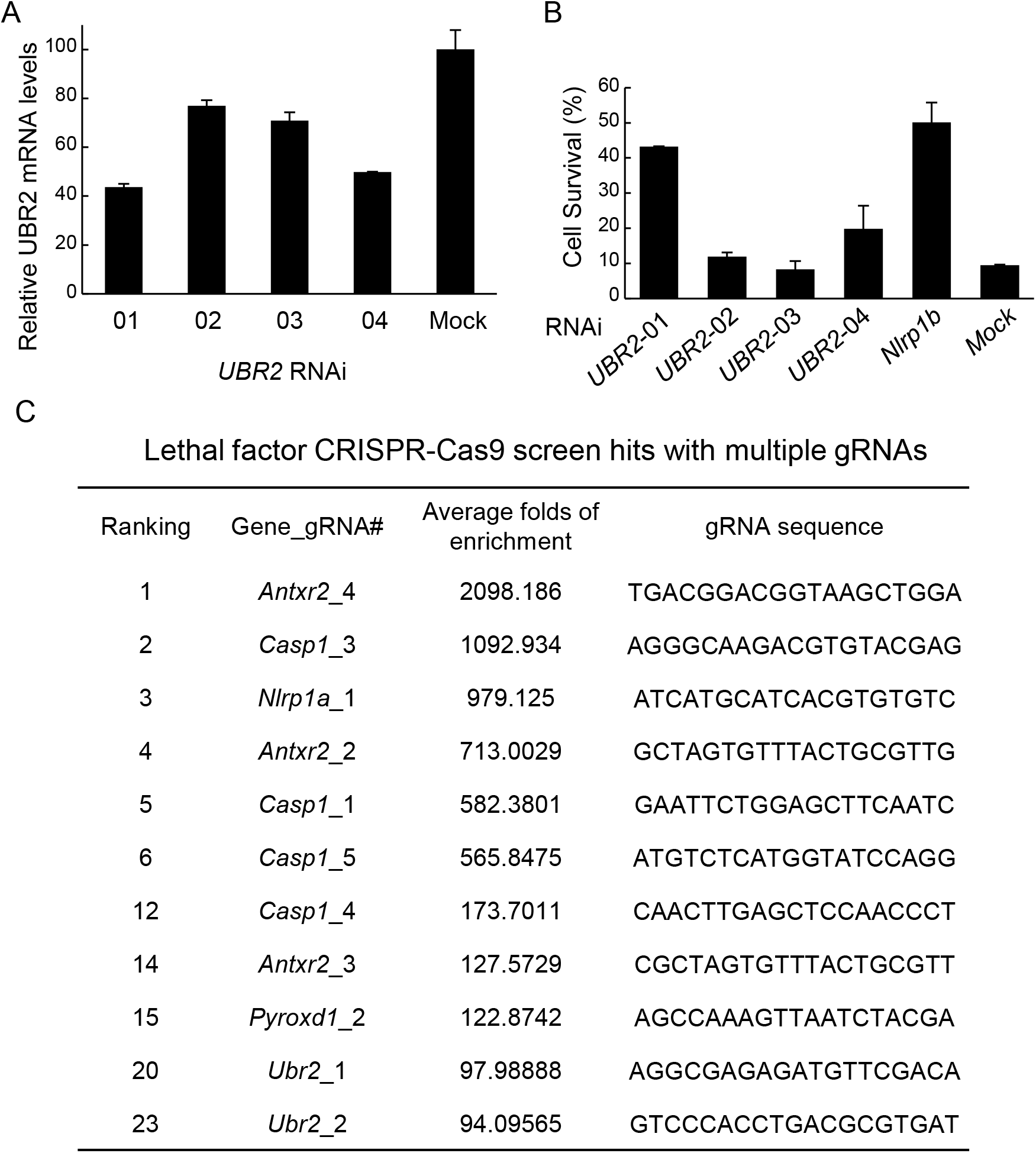
RNAi and CRISPR-Cas9 screen identified UBR2 in LF-induced Nlrp1b activation. (A) UBR2 knockdown efficiency measured by qPCR. MOCK, scramble siRNA. (B) Percentage of cell viability in UBR2-knockdown RAW^RA^ cells after LF treatment. RAW^RA^ cells were transfected with indicated siRNA for 60 h followed by LF treatment. Cell viability was measured by ATP cell viability assay. (C) gRNA hits from a genome-wide CRISPR-Cas9 screen of LF-induced pyroptosis in mouse Tlr4^-/-^ iBMDM cells. Shown are those genes with multiple gRNA hits. The ranking, the average fold increase and the sequences for each gRNA are listed. Note: The mouse gRNA library are designed for C57 BL/6J strain. The Nlrp1a_1 gRNA in the library match the Nlrp1b gene of the iBMDM cells we use for screen. But the four out of the five Nlrp1b gRNA in the library doesn’t match the Nlrp1b gene of the iBMDM cells, which is derived from 129/Sv mouse.

**Figure S4.**
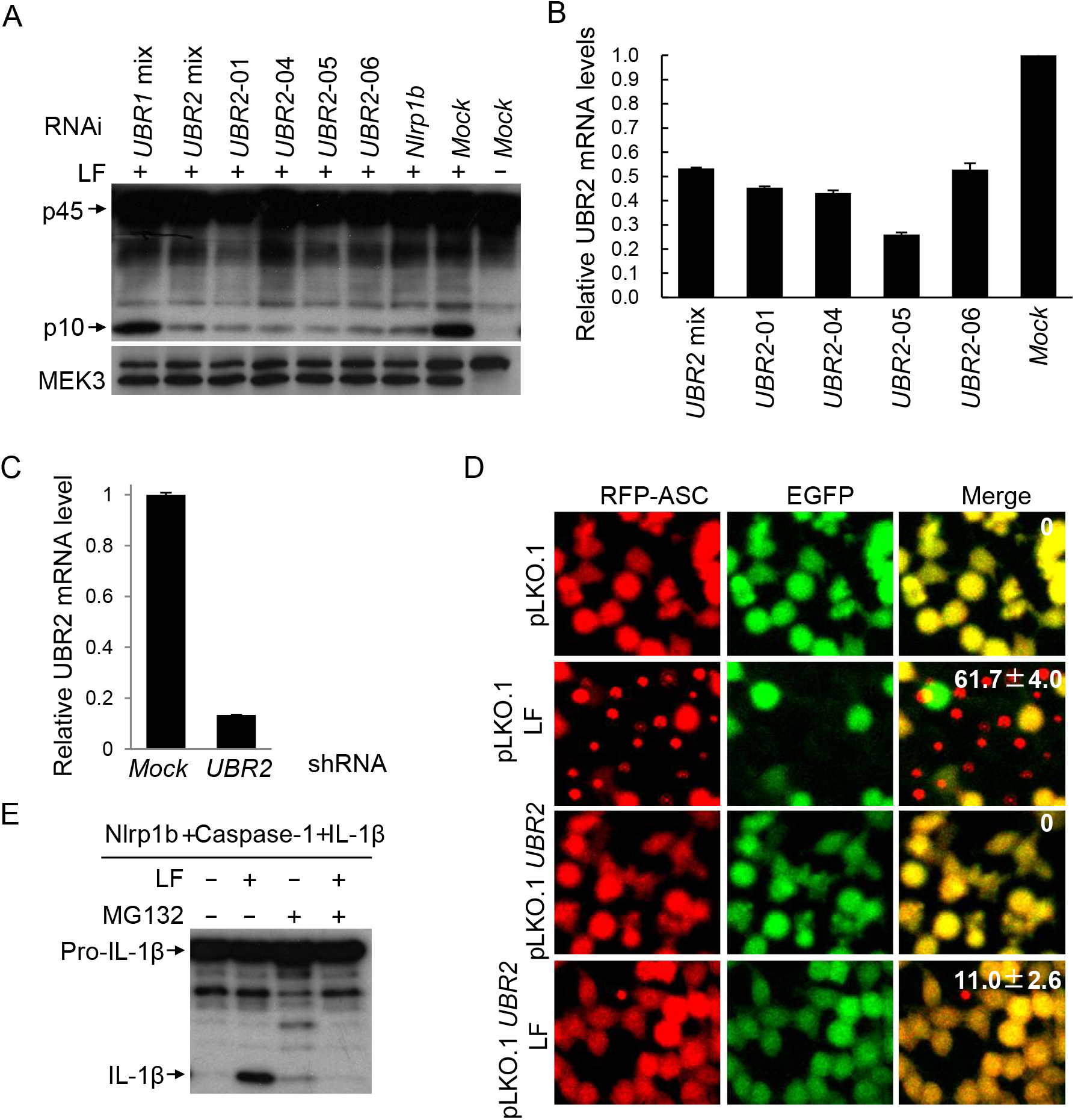
The N-end Rule E3 ligase UBR2 is required for LF-mediated Nalp1b activation. (A) Effects of UBR1, UBR2 or Nlrp1b knockdown on caspase-1 activation. Endogenous Nlrp1b or UBRs was silenced for 60 h before LF treatment and p10 western blotting. +, WT LF protein; –, LF E687C mutant protein. MOCK, scramble siRNA. (B) UBR2 knockdown efficiency in (A) measured by qPCR. MOCK, scramble siRNA. (C) UBR2 stable knockdown efficiency measured by qPCR. (D) RFP-ASC specks formation in RAW^RA^ cells with stable knockdown of UBR2. Statistics means percentage of RAW^RA^ cells in which RFP-ASC aggregates. (E) Reconstitution of Nalp1b inflammasome in 293T cells. 129/Sv mouse derived Nalp1b, human procaspase-1 and human pro-IL-1β expression plasmids were cotransfected in 293T cells, followed by induction with lethal toxin in the presence of 25 μM MG132 or not. +, WT LF protein; –, LF E687C mutant protein.

**Figure S5.**
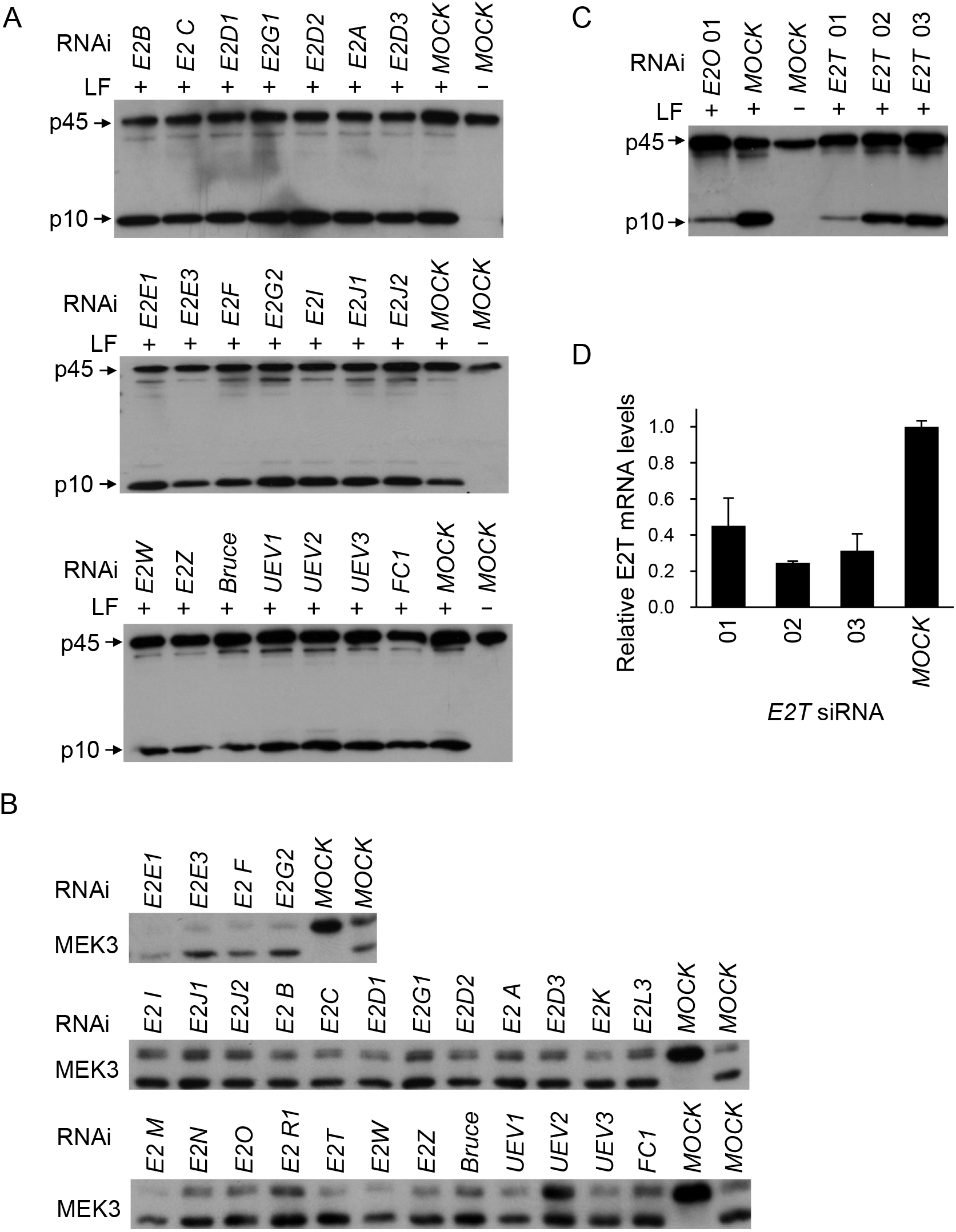
E2O participates in LF-triggered Nlrp1b activation. (A) Caspase-1 activation in E2s-knockdown 129/Sv mouse BMDM cells. BMDMs were transfected with indicated siRNA for 60 h, followed by induction with lf(+) or E687C mutant (−) protein. Caspase-1 activation was validated by p10 immunoblotting. MOCK, scramble siRNA. (B) MEK3 cleavage in E2s silencing BMDM cells. Cell lysates were made using sample derived from (A) and Figure 3D. (C) Additional E2T siRNAs were used to confirm the inhibitory effects of E2T knockdown on caspase-1 activation. BMDM cells were transfected with indicated siRNA respectively and then induced by LF. MOCK, scramble siRNA. (D) E2T knockdown efficiency of samples in (C)measured by qPCR.

**Figure S6.**
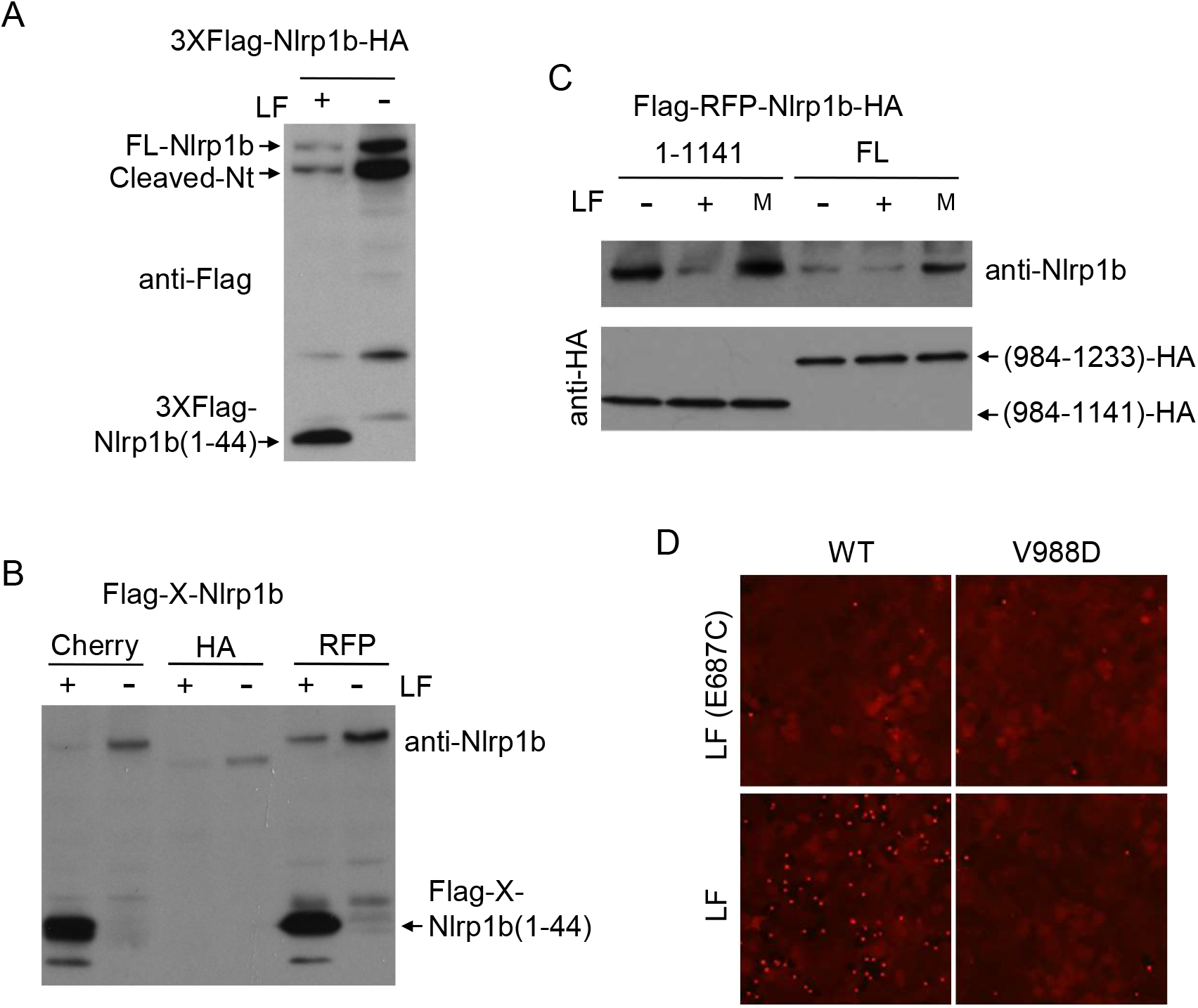
LF treatment induces the degradation of Nlrp1b. (A) Nlrp1b expression after LF treatment. 293T cells were transfected with tagged Nlrp1b plasmid for 24h, followed by induction with LF (+) or E687C mutant (−) protein. Nlrp1b and cleavage products after LF treatment were detected by Flag antibody. (B) Nlrp1b expression after LF treatment. 293T cells were transfected with indicated plasmids for 24 h, followed by induction with LF (+) or E687C mutant (−) protein. Nlrp1b expression was detected by Nlrp1b antibody. (C) Full-length and truncated Nlrp1b n expression after LF treatment. 293T cells were transfected with indicated plasmids for 24 h, followed by induction with no treat (−), LF (+) or E687C mutant (M) protein treatment. Nlrp1b expression was detected by Nlrp1b antibody. The C-terminal auto-cleavage products were detected by HA antibody. (D) RFP-ASC specks formation in RAW^RA^ cells expressing Nlrp1b WT and V988D mutant. RAW^RA^ cells were transfected with indicated plasmids for 24 h, followed by treatment with LF (+) or E687C mutant (−) protein.

**Table 1.**
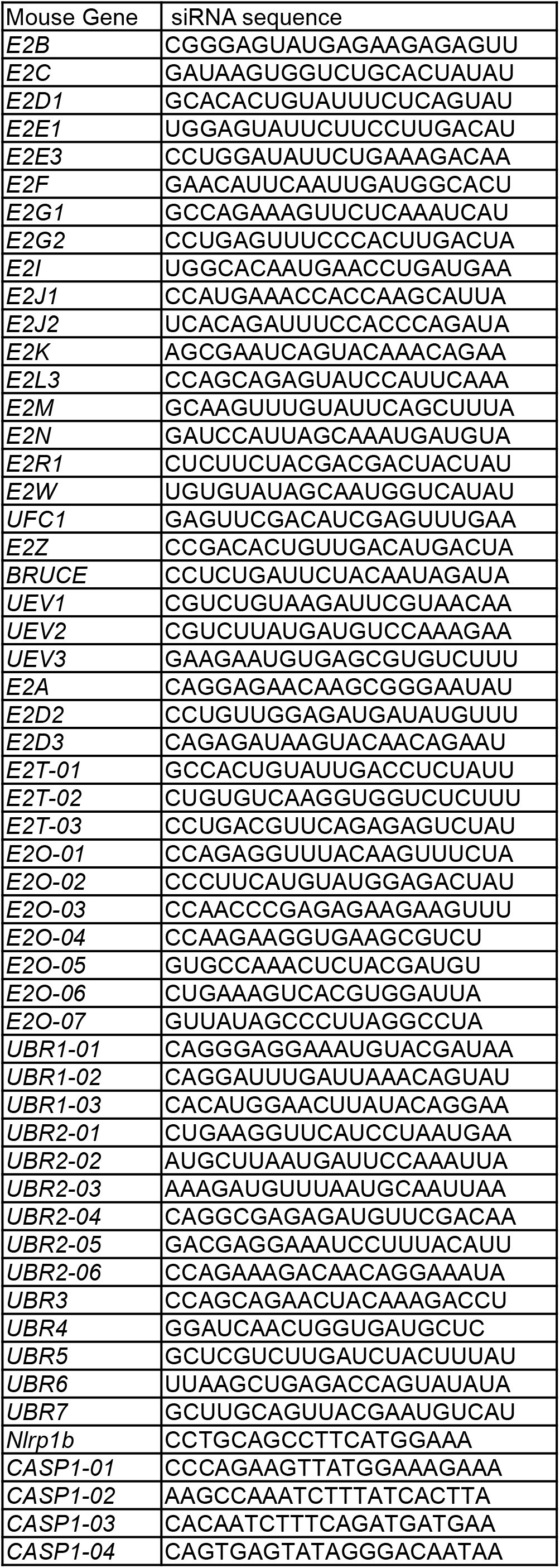
siRNA sequence.

**(S1).**
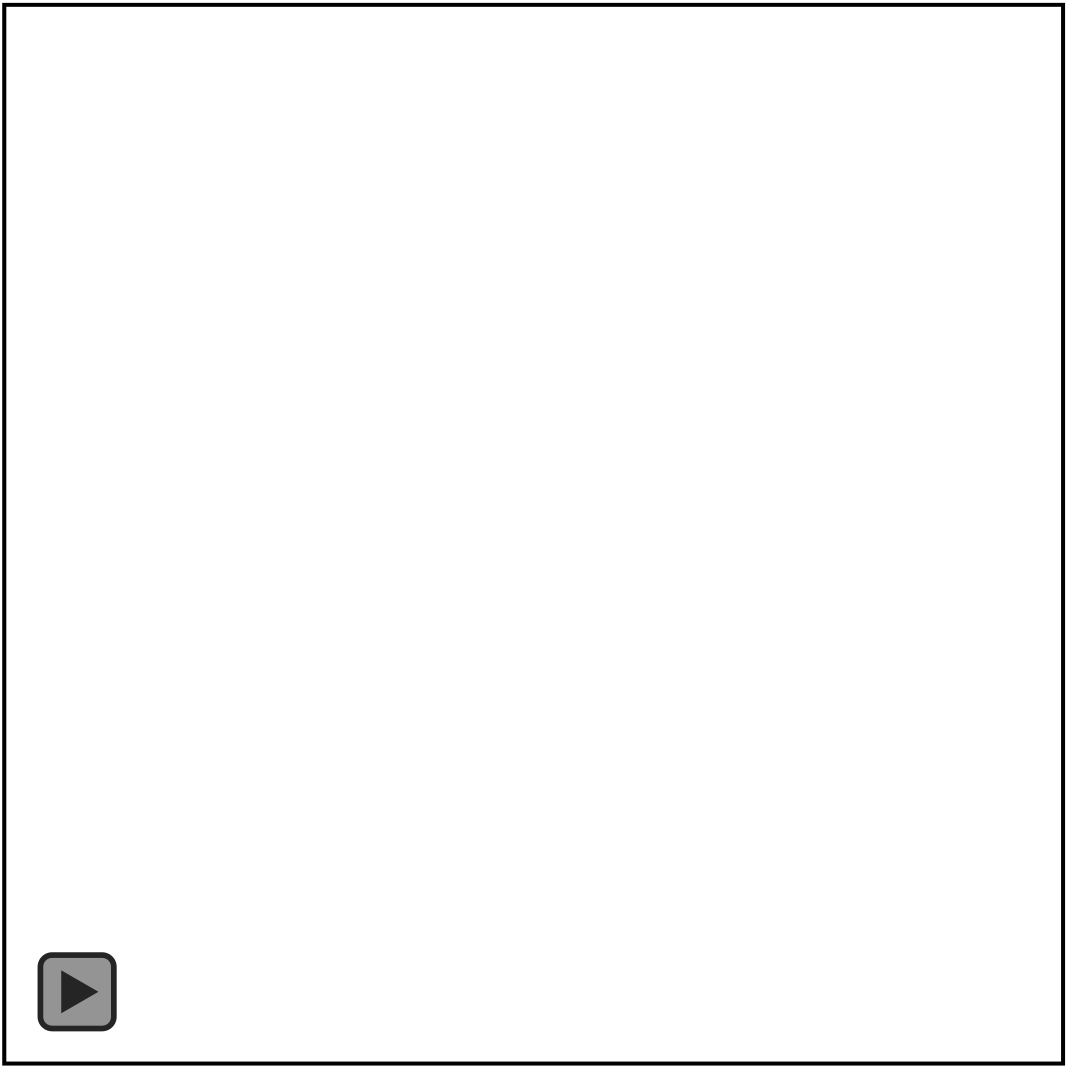
Movies RAW^RA^ cells morphology changes induced by LF. RAW^RA^ cell plated in 96-well format were incubated with LF plus protective antigen protein. DIC, EGFP and RFP images were taken using PerkinElmer Ultra-View spinning disc confocal.

**(S2).**
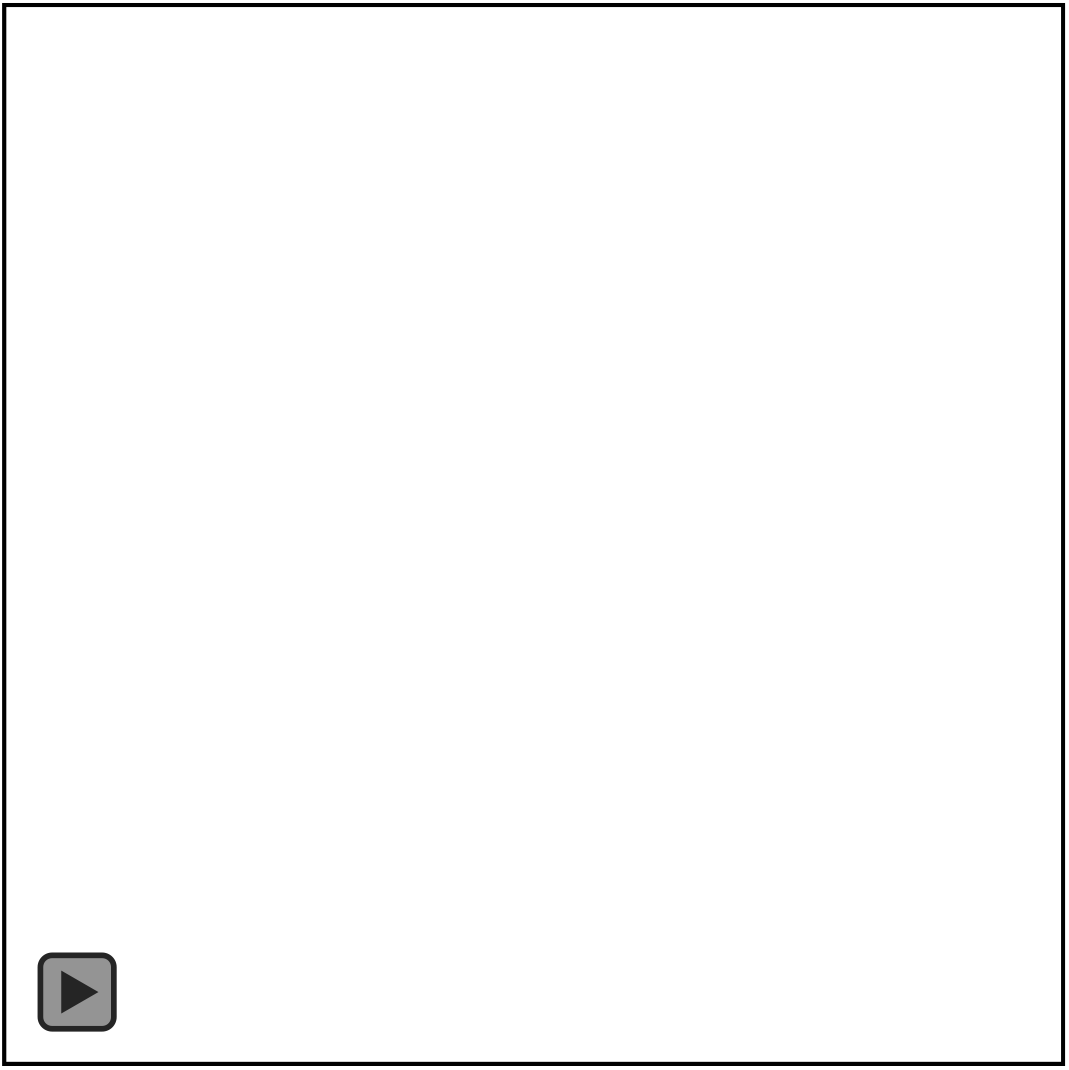
RAW^RA^ cell plated in 96-well format were incubated with LF E687C mutant plus protective antigen protein. DIC, EGFP and RFP images were taken using PerkinElmer Ultra-View spinning disc confocal.

